# Tenascin N contributes to spinal motor nerve morphogenesis during development

**DOI:** 10.64898/2026.01.29.702601

**Authors:** Charles G. Marcucci, Marieke Jones, Coleman Blanton, Sarah Kucenas

**Author notes:** **Contact Information**: Sarah Kucenas, Department of Biology, PO Box 400328, University of Virginia, Charlottesville, VA 22904-4328, USA.

## Abstract

Spinal motor nerves are an integral component of the nervous system whose development requires the coordination of many diverse cell types, including motor neurons, glia, and muscle. Although several molecular mechanisms guiding these interactions are known, many remain to be uncovered. Extracellular matrix (ECM) proteins also play a critical role in motor nerve assembly, yet their functions are less understood compared to classical pathfinding and guidance cues. Here, we identify a role for *tenascin-n* (*tnn*), an ECM glycoprotein, in spinal motor nerve development in zebrafish. Using *in situ* hybridization and immunohistochemistry, we show that *tnn/*Tnn is expressed and localized along vertical myosepta and the border of the ventral neural tube during spinal motor nerve development. To assess its function, we generated a CRISPR/Cas9 mutant allele, *tnn^uva96^*, and performed *in vivo* imaging and morphological analysis throughout motor nerve development. Loss of *tnn* leads to a subtle and transient increase in ectopic motor axon exit and aberrant motor axon branching in the zebrafish trunk. Our findings reveal a previously unrecognized role for *tnn* in spinal motor nerve assembly and expand our understanding of the diverse molecular contributors to spinal motor nerve development and morphogenesis.

## Introduction

Vertebrate spinal motor nerve development is a tightly orchestrated process that involves coordinated construction by multiple cell types, including neurons, the myotome, and glial cells. Many studies have identified factors crucial for their assembly, including but not limited to: intrinsic factors, mediators of cell-cell interactions, molecular gradients, and extracellular matrix proteins including collagens, chondroitin sulfate proteoglycans, and Tenascin-C (Guillon et al., 2016; Hilario et al., 2010; Schneider and Granato, 2006; Bernhardt and Schachner, 2000; Schweitzer et al., 2005; Nemoz-Billet et al., 2024). However, we have yet to uncover all the molecular mechanisms involved and how they cooperate to coordinate motor nerve assembly.

All vertebrates undergo a similar process of spinal motor nerve assembly that involves pioneering motor axons, molecular guidance of a leading growth cone, association of glial cells, and innervation of muscle targets (Bonanomi and Pfaff, 2010; Clotman and Tissir, 2020). Numerous vertebrate models have provided important insights into elucidating the molecular mechanisms that control these processes, including zebrafish (*Danio rerio*). Zebrafish are an excellent and often used model to study the developmental dynamics of spinal motor nerves as the sequence of events is conserved across vertebrates with the added benefit that they can be easily genetically manipulated and imaged *in vivo*. In this model, motor nerve assembly begins at approximately 18 hours post fertilization (hpf) as the three primary motor neurons (MNs) [called rostral primary (RiP), middle primary (MiP), and caudal primary (CaP)] extend their axons out of the ventral neural tube at sites called motor exit point transition zones (MEP TZs) (Myers, 1985; Myers et al., 1986; Westerfield et al., 1986). Simultaneously, neural crest cells begin streaming ventrally toward exiting motor axons (Raible et al., 1992; Schilling and Kimmel, 1994). Following axon extension by the three primary MNs, secondary MNs extend axons into the periphery at approximately 22 hpf, increasing in number over time (Pike et al., 1992; Beattie et al., 1997). Between 40 and 72 hpf, motor exit point (MEP) glia and perineurial glia (PG) exit the neural tube and associate with spinal motor nerves, where they will eventually myelinate proximal axon segments and ensheath nerve fascicles, respectively (Kucenas et al., 2008; Binari et al., 2013; Smith et al., 2014, Morris et al., 2017). Around this same stage, neural crest cells associated with nascent motor nerves begin differentiating into Schwann cells and myelinate peripheral motor axons (Raible and Eisen, 1994; Schilling and Kimmel, 1994; Brösamle and Halpern, 2002; Lyons et al., 2005). By 96 hpf, zebrafish larvae have fully functional peripheral motor nerves, including myelinated axons and nerve fascicle ensheathment by PG. In mice, these same events occur. However, the process takes multiple weeks beginning with motor neuron specification and axon extension at E10.5 with the achievement of adult-like movement by P21 (Jessen & Mirsky, 2015; Bucchia et al., 2018; Patel et al., 2022).

Many molecules are critical for guiding motor axons out of the central nervous system (CNS) and directing them to their muscle targets in the periphery, with diffusible attractant or repellant cues and their cognate receptors being the most classically studied, including Slit/Robo, Netrin-1/DCC, Semaphorins/Plexins, and Cxcl12/Cxcr4 (Rothberg et al., 1990; Brose et al., 1999; Huber et al., 2005; Lieberam et al., 2005; Palaisa & Granato, 2007; Kim et al., 2015; Kim et al., 2017;). Less often studied, however, are extracellular matrix (ECM) proteins. A few with known roles include collagens, chondroitin-sulfate proteoglycans (CSPGs), and Tenascin-C (Guillon et al., 2016; Hilario et al., 2010; Schneider and Granato, 2006; Bernhardt and Schachner, 2000; Schweitzer et al., 2005; Nemoz-Billet et al., 2024). Multiple collagen proteins, including ones encoded by *col19a1*, *col18a1*, and *col15a1b* are crucial for directing peripheral motor axons to their muscle target. For example, loss of *col15a1b* results in defects in primary and secondary motor axon pathfinding and subsequent muscle atrophy (Guillon et al., 2016; Hilario et al., 2010; Schneider and Granato, 2006, Nemoz-Billet et al., 2024). Removal of CSPGs via treatment with the enzyme chondroitinase ABC results in aberrant branching of ventral motor axons (Bernhardt and Schachner, 2000). Finally, loss of *tenascin-C* (*tnc*) results in aberrant branching of motor axons at a developmental choice point of growing motor axons called the horizontal myoseptum at 33 hpf and increased branching of ventral motor axons at 27 hpf in zebrafish (Schweitzer et al., 2005; Nemoz-Billet et al., 2024). Constituents of the extracellular matrix are clearly important contributors to spinal motor nerve development, yet we still have not uncovered all the key players nor elucidated their roles in spinal motor nerve assembly.

In our studies, we sought to investigate whether an understudied tenascin family member, Tenascin-N (Tnn), also played a role in spinal motor nerve development. Tnn, originally named Tnw, was first described by Philipp Weber and colleagues in zebrafish as the fourth member of the tenascin family, and later named Tnn after the homolog was discovered and named in mice (Weber et al., 1998; Neidhardt et al., 2003). In their initial description, Weber and colleagues described partially overlapping expression of *tnn* and *tnc* in the developing zebrafish embryo (Weber et al., 1998). The initial study in which Tnn was identified in mice demonstrated that it repelled neurite growth of hippocampal cells in culture (Neidhardt et al., 2003). Additionally, Tnn is implicated in stem cell niches, cancer cell signaling, and tumor cell migration and metastasis (Tucker and Degen, 2019 review). However, there are no studies investigating Tnn in spinal motor nerve development.

In this study, we identify a transient role for Tnn in spinal motor development. Expression analysis revealed that *tnn* is expressed between 1 and 3 days post fertilization (dpf) in zebrafish embryos and larvae, a critical window during spinal motor nerve assembly. To investigate its function, we generated a null mutant allele using CRISPR/Cas9 genome editing and characterized this line through immunohistochemistry and *in vivo* imaging. Loss of *tnn* resulted in a transient higher incidence of ectopic spinal motor nerve exits at 2 dpf and increased motor axon arborization at 2 and 3 dpf. These findings demonstrate that *tnn* contributes to early spinal motor nerve morphogenesis. By focusing on understudied ECM component, our work reveals a previously unrecognized contributor to motor nerve development and underscores the need to further explore the role of ECM in spinal motor nerve assembly.

## Results

### *Tnn/*TNN is expressed along vertical myosepta and the ventral border of the spinal cord during zebrafish development

To begin investigating the role of Tnn in spinal motor nerve development, we first sought to investigate its expression pattern in the developing zebrafish embryo between 1 and 3 dpf. We examined both RNA and protein expression, and our data confirmed initial findings reported by Weber et al. (Weber et al., 1998). In whole mount preparations from 1 to 3 dpf, we observed *tnn* and Tnn expression along vertical myosepta bordering each somite (analogous to myotendinous junctions in mammals) (Figure 1A-F’). At 2 and 3 dpf but not 1 dpf, we also observed Tnn localization adjacent to the ventral boundary of the neural tube, including at MEP TZs, indicated by staining of motor axons with an anti-acetylated tubulin antibody (Figure 1B, B’, D, D’, F, F’). To better assess the localization of this expression, we cryosectioned 2 and 3 dpf larvae and performed *in situ* hybridization (chromogenic and fluorescent) and immunohistochemistry on transverse sections of the trunk. In these sections we observed *tnn* expression bordering the ventral neural tube, including adjacent to MEP TZs indicated by peripheral *nkx2.2a* expression driven by the *Tg(nkx2.2a:megfp)* transgenic line (Figure 1G-H’) as well as Tnn localization along vertical myosepta (Figure 1H’’). From our data, we conclude that *tnn*/Tnn is localized along vertical myosepta and the ventral neural tube, including at MEP TZs.

**Figure 1.**
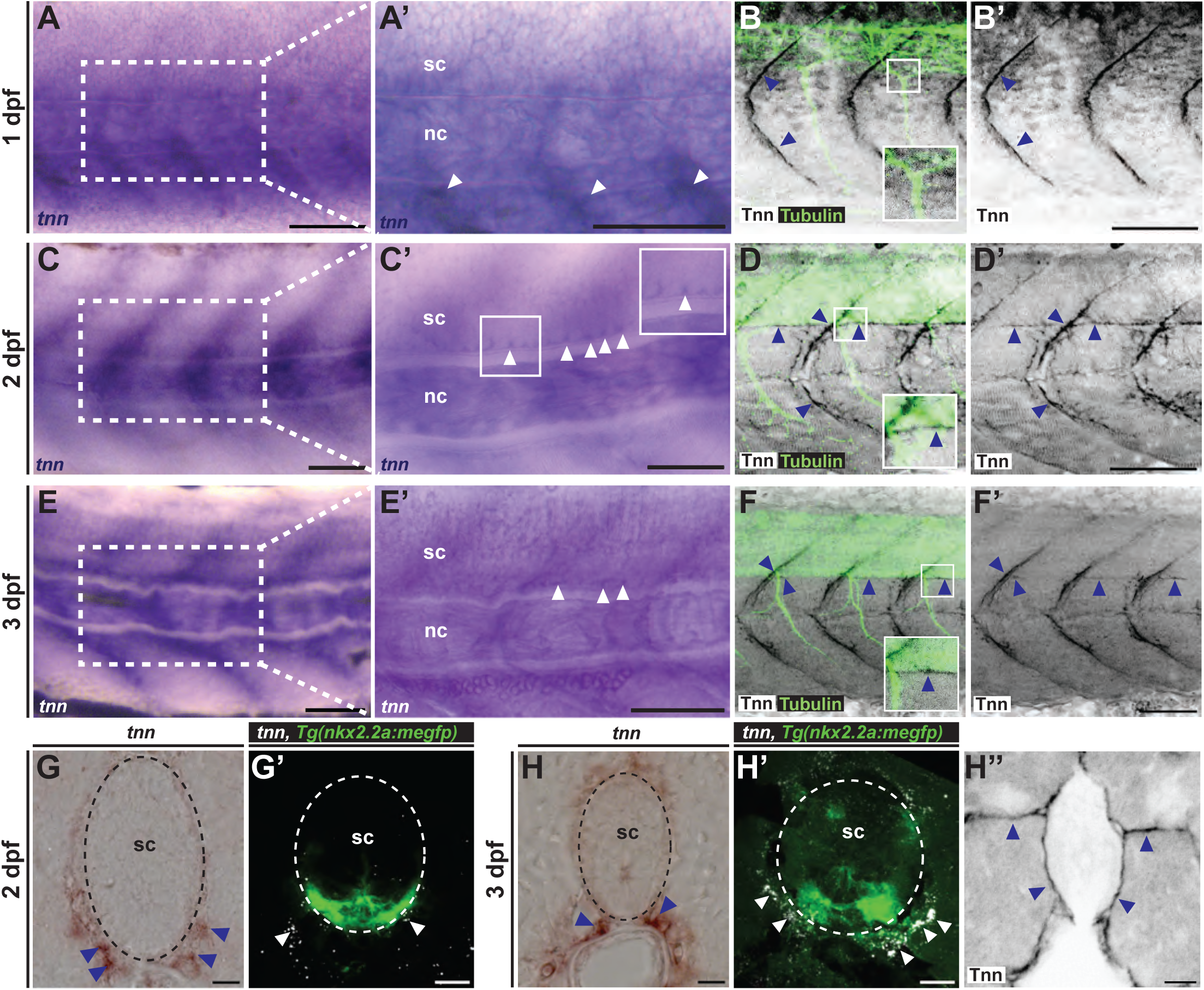
*Tnn/*TNN is expressed along vertical myosepta and the ventral border of the spinal cord during zebrafish development. (**A,C,E**) Bright-field images from a lateral view of whole mount chromogenic *in situ* hybridization (ISH) with enlarged images (**A’, C’, E’**) showing *tnn* expression (purple) along vertical myosepta and adjacent to the ventral neural tube (white arrowheads in enlarged images) at 1, 2, and 3 days post fertilization (dpf). (**B-B’, D-D’, F-F’**) Immunofluorescence showing Tnn (black) and Tubulin (green) localized along vertical myosepta and ventral neural tube (blue arrowheads) at 1, 2, and 3 dpf (**B-B’, D-D’,** and **F-F’** respectively). Insets in **D** and **F** show Tnn at sites of motor axon exit at 2 and 3 dpf, respectively. (**G**) Bright-field image of chromogenic ISH in transverse section of the trunk showing *tnn* expression (brown) in cells adjacent to the ventral neural tube (blue arrowheads) at 2 dpf. (**G’**) Fluorescent ISH showing *tnn* expression (white) adjacent to the ventral neural tube (white arrowheads) at 2 dpf in transverse section of *Tg(nkx2.2a:megfp)* larvae. (**H**) Bright-field image of chromogenic ISH showing *tnn* expression (brown) in cells adjacent to the ventral neural tube (blue arrowheads) at 3 dpf in transverse section. (**H’**) Fluorescent ISH showing *tnn* expression (white) adjacent to the ventral neural tube (white arrowheads) at 3 dpf in transverse section of *Tg(nkx2.2a:megfp)* larvae. (**H’’**) Immunofluorescence showing Tnn (black) bordering the neural tube and along vertical myosepta (blue arrowheads). All images oriented with anterior to the left, dorsal to the top. Scale bars: (**A-F**) 50 μm; (**G-H**) 20 μm. sc = spinal cord; nc = notochord.

### Generation of *tnn* null mutant allele with CRISPR/Cas9 genome editing

After observing that *tnn* was expressed in a pattern that could influence spinal motor nerve growth, we hypothesized that it may contribute to development of these structures. To investigate this hypothesis, we created genetic mutants using CRISPR/Cas9 genome editing (Hwang et al., 2013; Gagnon et al., 2014). We designed a guide RNA (gRNA) to target the sixth coding exon of *tnn* and injected the gRNA with Cas9 protein into wild-type embryos at the one-cell stage (Figure 2A-B). After raising F0 embryos to adulthood, we identified a founder, *tnn^uva96^,* which harbors a four base pair insertion in the sixth coding exon that codes for the third of five-fibronectin type III (FNIII) domains (Figure 2B-C). This insertion results in a missense (W502L) and nonsense mutation at amino acid positions 502 and 503, respectively, truncating the sequence by 429 amino acids, eliminating two FNIII domains and the most C-terminal domain, a fibrinogen-related domain (Figure 2C-D).

**Figure 2.**
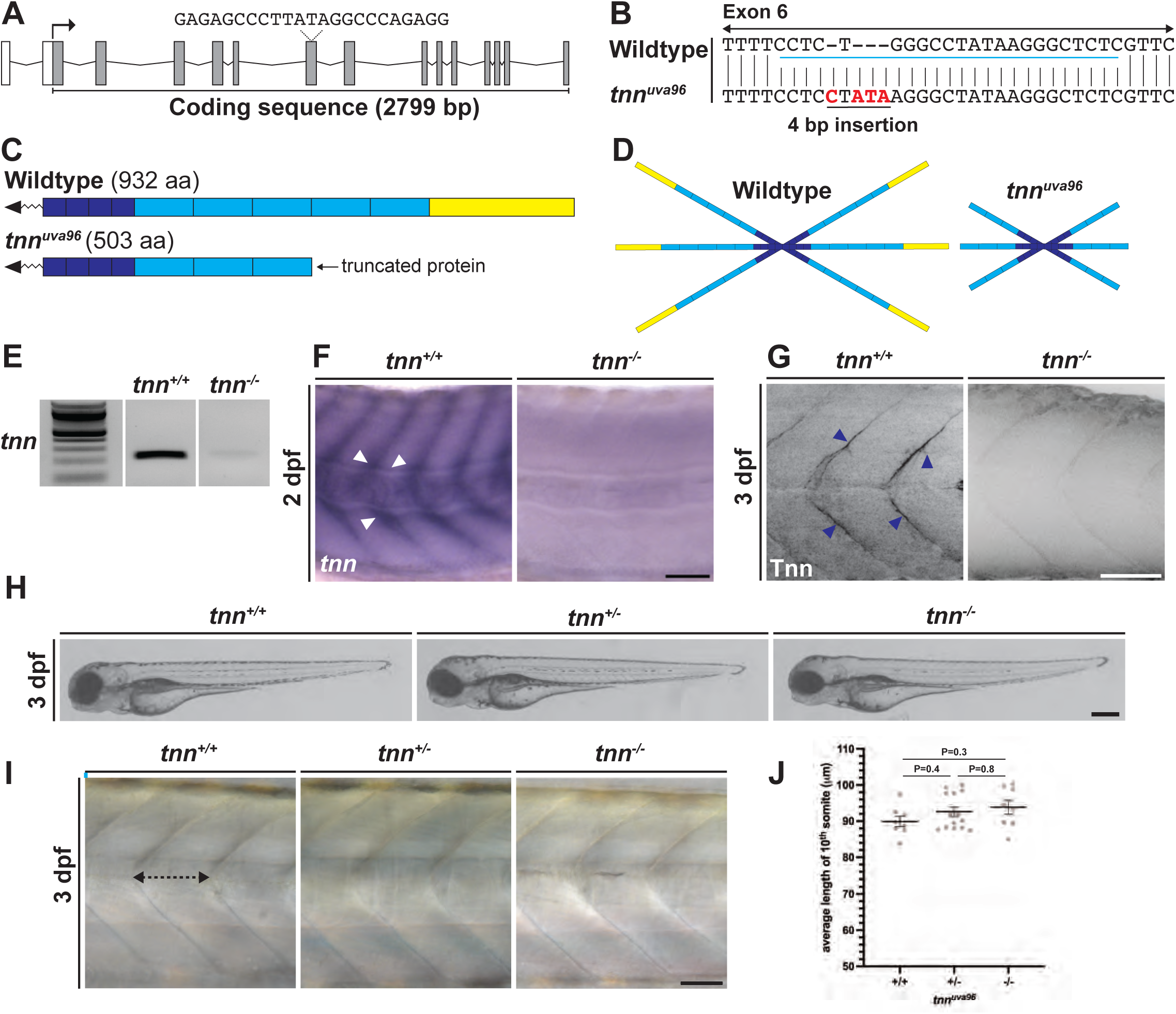
Generation of *tnn* null mutant allele using CRISPR/Cas9 genome editing. (**A**) Schematic of zebrafish *tnn* gene. The coding sequence (grey; 2799 bp) encodes the protein domains represented in (**C**), and the non-coding sequence is represented in white. Dashed lines above coding exon 6 in schematic indicate the gRNA sequence and target region of the coding sequence. (**B**) Genomic sequences for wild-type *tnn* and and mutant *tnn^uva96^*showing a 4 bp insertion within exon 6 of the *tnn* gene. (**C**) Schematic of Tnn protein made in wild-type (932 aa; top) and predicted protein made in *tnn^uva96^* mutant fish (503 aa; bottom; dark blue: epidermal growth factor-like domains, light blue: fibronectin type III domains, yellow: fibrinogen-related domain). (**D**) Schematic showing that wild-type Tnn assembles into a hexabrachion made up of two Tnn trimers with a predicted hexabrachion structure in *tnn^uva96^* mutants. (**E**) Gel electrophoresis image showing wild-type and *tnn^uva96^* RT-PCR products at 2 dpf. (**F**) Bright-field images of chromogenic *in situ* hybridization (ISH) showing *tnn* (purple) in wild-type larvae along vertical myosepta and ventral neural tube (white arrowheads) and the absence of *tnn* expression in *tnn^uva96^* homozygous larvae at 2 dpf. (**G**) Immunofluorescence showing Tnn (black) localized along vertical myosepta and ventral neural tube (blue arrowheads) in wild-type larvae and the absence of Tnn antibody labeling in *tnn^uva96^* homozygous mutant larvae at 3 dpf. (**H**) Brightfield images of wild-type (left), *tnn^uva96^*heterozygous (middle), and *tnn^uva96^* homozygous (right) larvae at 3 dpf showing no gross anatomical defects in *tnn^uva96^* mutants. (**I**) Differential interference contrast (DIC) images at the level of the 10^th^ somite in wild-type, *tnn^uva96^* heterozygous, and *tnn^uva96^* homozygous mutant larvae at 3 dpf. Black dotted line capped with arrowheads represents the measurements taken in each image for quantification in (**J**). (**J**) Scatter plot of the length of the 10^th^ somite at 3 dpf (dot = measurement from 1 larva, wild-type n=8 heterozygous n=14, homozygous n=8). Data compared with a one-way ANOVA with Tukey’s post-hoc test. All images oriented with anterior to the left, dorsal to the top. Scale bars: (**F,G,H**) 50 μm; (**I**) 0.25 mm;

To determine if this mutation resulted in a null allele, we performed RT-PCR, whole mount RNA *in situ* hybridization, and immunohistochemistry. Using all three methods, we observed no *tnn/*Tnn expression in homozygous mutant larvae at either 2 or 3 dpf (Figure 2E-G). From these data we conclude that the *tnn^uva96^* allele is a null allele which provides us with a line with which to interrogate the functional role of *tnn* in spinal motor nerve assembly.

To begin characterizing *tnn^uva96^* mutant embryos and larvae, we analyzed their general development and morphology. At 3 dpf, we did not observe any gross morphological abnormalities between wild-type, heterozygous, and mutant larvae (Figure 2H). We also measured somite width at the level of the 10^th^ somite and saw no significant difference between the genotypes (Figure 2I-J). Additionally, both heterozygous and homozygous mutant animals survived to adulthood and successfully reproduced.

### *tnn* contributes to motor axon exit and branching of spinal motor nerves at 2 dpf

Using our newly created mutant allele, *tnn^uva96^*, we investigated the role of *tnn* during spinal motor nerve development. To do this, we used *in vivo,* confocal imaging in *Tg(olig2:dsred)* larvae where *olig2* regulatory sequences drive expression of DsRed in motor neurons and axons. In wild-type larvae at 2 dpf, we observed uniform, discrete spacing of motor nerve exits along the anterior-posterior axis of the neural tube (Figure 3A-C). In contrast, in *tnn^uva96^* heterozygous and homozygous larvae, we observed an increased incidence of ectopic motor nerve exit compared to wild-type larvae (Figure 3A-C). Of all nerves imaged between somites 10 and 22, 6.3% and 5.7% were the result of ectopic exit in heterozygous and homozygous mutant embryos, respectively, compared to 3.0% in wild-type larvae (Figure 3B; Fisher’s exact test, P=0.3). Further, 50% and 52% of heterozygous and homozygous mutant larvae, respectively, displayed at least one ectopic motor nerve exit as compared to 31% of wild-type larvae (Figure 3C; Fisher’s exact test, P=0.4). To further quantify this disorganization, we measured the periodicity, or distance between each nerve exit and its nearest neighbor at 2 dpf. Heterozygous and homozygous mutant larvae had a slight decrease (P=0.4 and P=0.8, respectively) in average periodicity per larvae compared to wild-type siblings [wild-type estimated marginal mean [95%confidence interval (CI)] (emm)=76.2 μm [72.8, 79.7], heterozygous emm [95% CI]=73.4 μm [71.1, 75.6], homozygous emm [95% CI]=74.6 μm [71.3, 77.8])], and an increase in average variance of periodicity (P=0.5 and P=0.3) compared to wild-type siblings (wild-type emm [95% CI]=36.3 [17.4, 77.0], heterozygous emm [95% CI]=59.3 [36.2, 97.1], homozygous emm [95% CI]=75.8, 37.1, 155.2) (Figure 3D-E).

**Figure 3.**
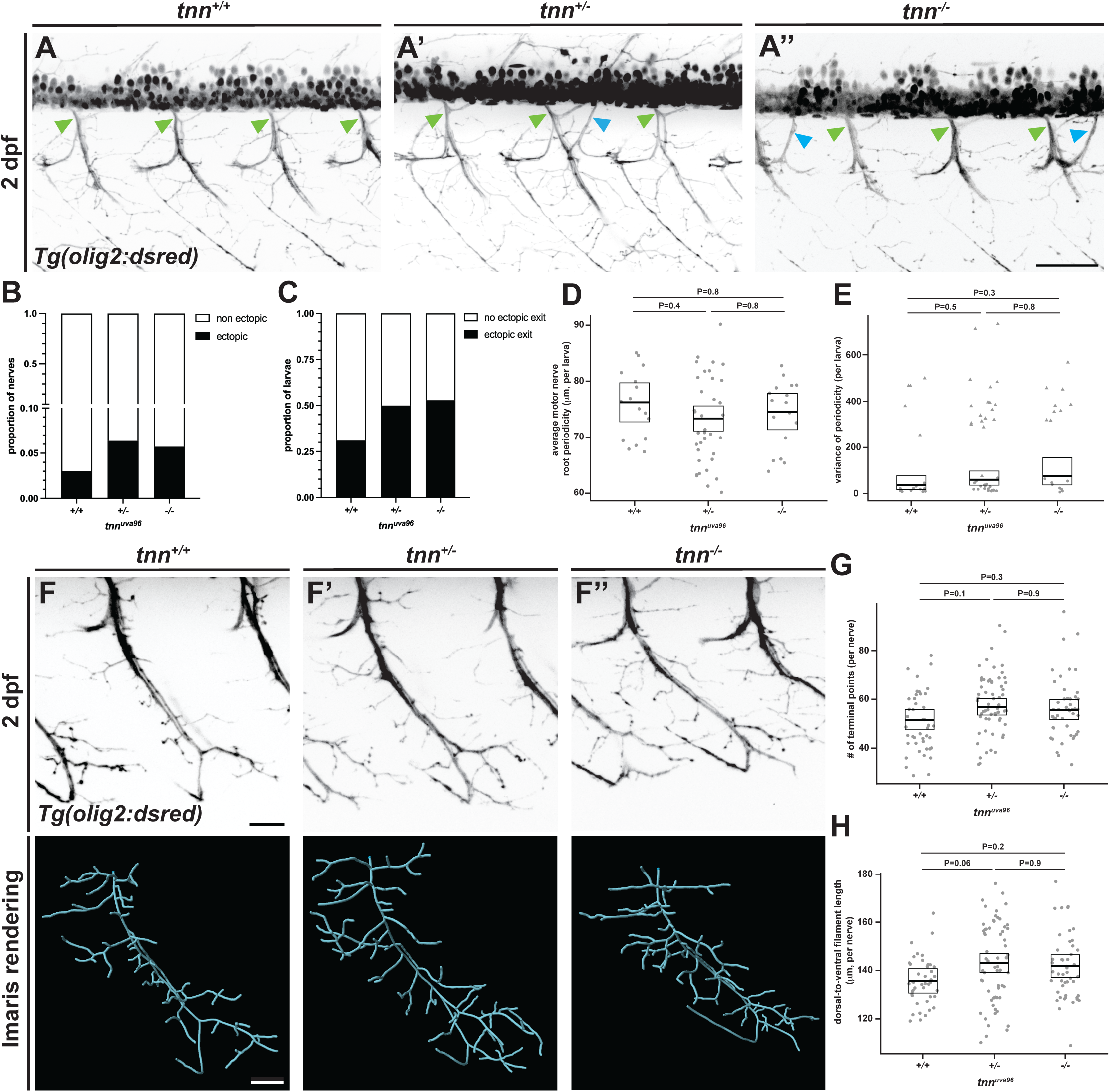
*tnn* contributes to motor axon exit and branching of spinal motor nerves at 2 dpf. (**A**) Representative images from *in vivo,* confocal images of laterally mounted *Tg(olig2:dsred)* wild-type (**A**), *tnn* heterozygous (**A’**), and *tnn* homozygous (**A’’**) embryos at 2 dpf showing instances of normally placed ventral motor nerve exits (green arrowheads) and ectopic motor nerve exits (blue arrowheads). (**B**) Stacked histogram of proportion of nerves that resulted from ectopic exit and non-ectopic exit (wild-type n=165, heterozygous n=408, homozygous n=192; Fisher’s exact test P=0.3). (**C**) Stacked histogram of proportion of larvae with at least one instance of ectopic nerve exit and larvae with no instances of ectopic exit (wild-type n=16, heterozygous n=36, homozygous n=17; Fisher’s exact test P=0.4). (**D**) Scatter plot of average periodicity per larva (measured as distance between each nerve exit and its nearest neighbor) (dot = 1 larva; wild-type n=16, heterozygous n=36, homozygous n=17). (**E**) Scatter plot of average variance of periodicity per larva (dot/triangle = 1 larva; triangle = larva with at least one instance of ectopic exit; wild-type n=16, heterozygous n=36, homozygous n=17). (**F**) Representative images (top images) from *in vivo,* confocal images of laterally mounted *Tg(olig2:dsred)* wild-type (**F**), *tnn* heterozygous (**F’**), and *tnn* homozygous (**F’’**) embryos at 2 dpf to show a single nerve selected for tracing with the corresponding Imaris 3D rendering produced from tracing shown below. (**G**) Scatter plot of the number of terminal points per nerve (dot = 1 nerve; wild-type n=43, heterozygous n=62, homozygous n=47). (**H**) Scatter plot of the distance from the most dorsal to most ventral point of each traced nerve (dot = 1 nerve; wild-type n=43, heterozygous n=62, homozygous n=47). Summary statistics for each scatter plot represent the estimated marginal mean (emm) with 95% confidence interval (CI) predicted from the described model. P-values generated from post-hoc Tukey HSD. All images oriented with anterior to the left, dorsal to the top. All images taken between somites 10 and 22. Scale bars: (**A’’**) 50 μm; (**F**) 20 μm.

In addition to perturbed motor nerve spacing at 2 dpf, we also observed irregularities in the branching of motor axons in mutant larvae. To thoroughly characterize this, we traced individual nerves using the Imaris imaging filament module (Figure 3F). From these traces, we used terminal points (a count of each branch ending) as a measurement of arborization. Heterozygous and homozygous mutant larvae had increased terminal points per nerve (P=0.1 and P=0.3) compared to wild-type larvae at 2 dpf (wild-type emm [95% CI]=51.5 [47.5, 55.8], heterozygous emm [95% CI]=56.7 [53.5, 60.2], homozygous emm [95% CI]=55.6 [51.7,59.8] terminal points) (Figure 3G-H).

To determine when ectopic exits occurred, we employed the same imaging strategy described above between 24 and 28 hours post fertilization (hpf) and observed initial breaches of ectopically exiting axons (Figure 4A). At this time point, of all nerves imaged, 3.6% and 3.9% were the result of ectopic exit in wild-type and heterozygous mutant larvae, respectively, compared to 0% in homozygous mutant larvae (Figure 4B; Fisher’s exact test, P=0.2). Further, 21.4% and 36.4% of wild-type and heterozygous mutant larvae, respectively, displayed at least one ectopic motor nerve exit as compared to 0% of homozygous larvae (Figure 4C; Fisher’s exact test, p=0.06). Additionally, there was a slight increase in average periodicity per larvae for homozygous mutant compared to wild-type and heterozygous mutant (P=0.6 and P=0.3) embryos, respectively, at 1 dpf (wild-type emm [95% CI]=59.9 μm [56.9, 63.0], heterozygous emm [95% CI]=59.1 μm [57.1, 61.1], homozygous emm [95% CI]=62.0 μm [58.2, 65.9]) (Figure 4D). We also observed a significant difference across all three genotypes in variance of periodicity (wild-type emm [95% confidence interval (CI)]=26.1 [25.9, 26.3], heterozygous emm [95% CI]=39.4 [39.0, 39.8], homozygous emm [95% CI]=13.0 [12.8, 13.1]; P<0.0001 for all comparisons) (Figure 4E). Additionally, using *in vivo,* time-lapse imaging starting between 24 and 28 hpf hours, we observed motor axons actively extend into the periphery at ectopic locations in all genotypes over the course of 16 hours (Supplemental Video 1). These findings align with the timing of Tnn localization as the appearance of Tnn along the ventral neural tube does not occur until sometime between 1 and 2 dpf (Figure 1B,D). Together, we conclude that Tnn influences the spacing of motor axon exit from the ventral neural tube and arborization of growing motor axons between 1 and 2 dpf.

**Figure 4.**
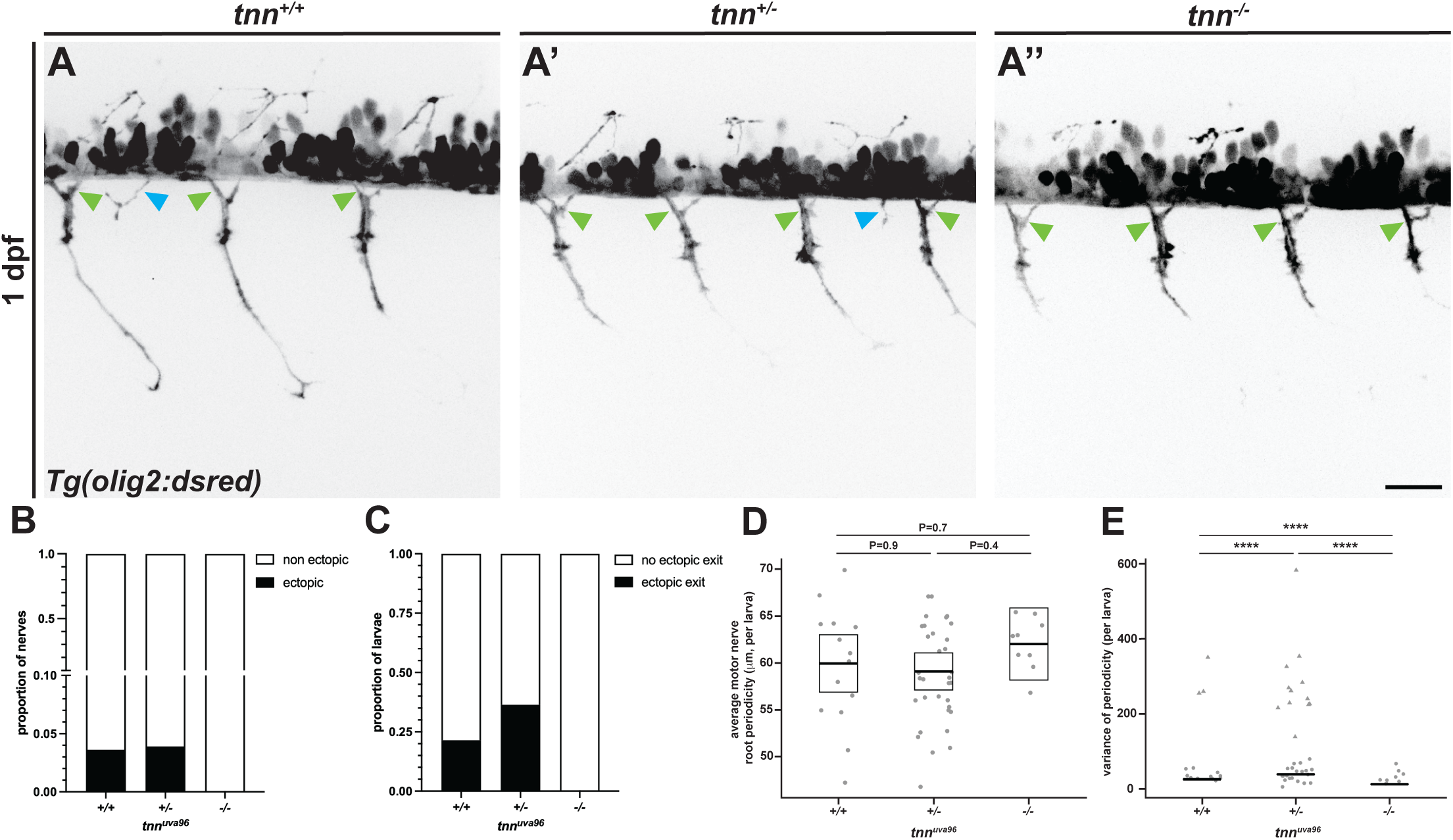
Ectopic exits appear at 1 dpf in *tnn^uva96^* larvae. (**A**) Representative images from *in vivo,* confocal images of laterally mounted *Tg(olig2:dsred)* wild-type (**A**), *tnn* heterozygous (**A’**), and *tnn* homozygous (**A’’**) embryos at 1 dpf [(between 24-28 hours post fertilization)] showing instances of normally placed ventral motor nerve exits (green arrowheads) and ectopic motor nerve exits (blue arrowheads). (**B**) Stacked histogram of proportion of nerves that resulted from ectopic exit and non-ectopic exit (wild-type n=138, heterozygous n=334, homozygous n=85; Fisher’s exact test P=0.2). (**C**) Stacked histogram of proportion of embryos with at least one instance of ectopic nerve exit and embryos with no instances of ectopic exit (wild-type n=14, heterozygous n=33, homozygous n=10; Fisher’s exact test P=0.058). (**D**) Scatter plot of average periodicity per embryo (measured as distance between each nerve exit and its nearest neighbor) (dot = 1 embryo, wild-type n=14, heterozygous n=33, homozygous n=10). (**E**) Scatter plot of average variance of periodicity per embryo (dot/triangle = 1 embryo; triangle = embryo with at least one instance of ectopic exit; wild-type n=14, heterozygous n=33, homozygous n=10). **** (P<0.0001). Summary statistics for each scatter plot represent the emm with 95% CI predicted from the described model. P-values generated from post-hoc Tukey HSD. Images oriented with anterior to the left, dorsal to the top. All images taken between somites 10 and 22. Scale bar: (**A’’**) 50 μm.

### Disruptions to peripheral spinal motor nerve exit and arborization are transient in *tnn^uva96^* homozygous larvae

To determine whether the disruptions to nerve exit and branching persisted as spinal motor nerve development progressed, we repeated *in vivo* imaging in 3 dpf *tnn^uva96^;Tg(olig2:dsred)* larvae (Figure 5A). Interestingly, of all nerves imaged, 6.9% were the result of ectopic exit in heterozygous mutant larvae, compared to 4.5% and 4.3% for wild-type and homozygous mutant larvae, respectively (Figure 5B; Fisher’s exact test, P=0.4). Further, 56% percent of heterozygous mutant larvae displayed at least one ectopic nerve exit, compared to 36% and 29% of wild-type and homozygous mutant larvae (Figure 5C; Fisher’s exact test, P=0.2). Examining periodicity at 3 dpf, we observed no significant difference between genotypes in average periodicity per larvae (wild-type emm [95% CI]=79.2 μm [75.2, 83.1], heterozygous emm [95% CI]=79.3 μm [76.5, 82.2], homozygous emm [95% CI]=80.5 μm [76.9, 84.1]) (Figure 5D). We did, however, observe significant differences in variance of periodicity across all three genotypes (wild-type emm [95% CI]=42.7 [42.3, 43.1], heterozygous emm [95% CI]=55.5 [54.8, 56.2], homozygous emm [95% CI]=37.7 [37.2, 38.2]; P<0.0001 for all comparisons) (Figure 5E). We also performed the same motor nerve tracing to characterize branching of ventral motor nerves at 3 dpf (Figure 5F). Heterozygous and homozygous mutant larvae showed slightly decreased motor axon arborization compared to wild-type larvae (P=0.2 and P=0.3) at 3 dpf (wild-type emm [95% CI]=109 [102.4, 117], heterozygous emm [95% CI]=102 [97.2, 107], homozygous emm [95% CI]=102 [95.8, 109] terminal points) (Figure 5G-H). Taken together, we did not observe the persistence of increased ectopic exit or motor axon arborization at 3 dpf in homozygous mutant larvae, while heterozygotes did continue to display an increased incidence of ectopic exit and a significant increase in variance of periodicity. These findings led us to hypothesize that genetic compensation induced by loss of two copies of *tnn* may be contributing to these differences seen at 2 versus 3 dpf.

**Figure 5.**
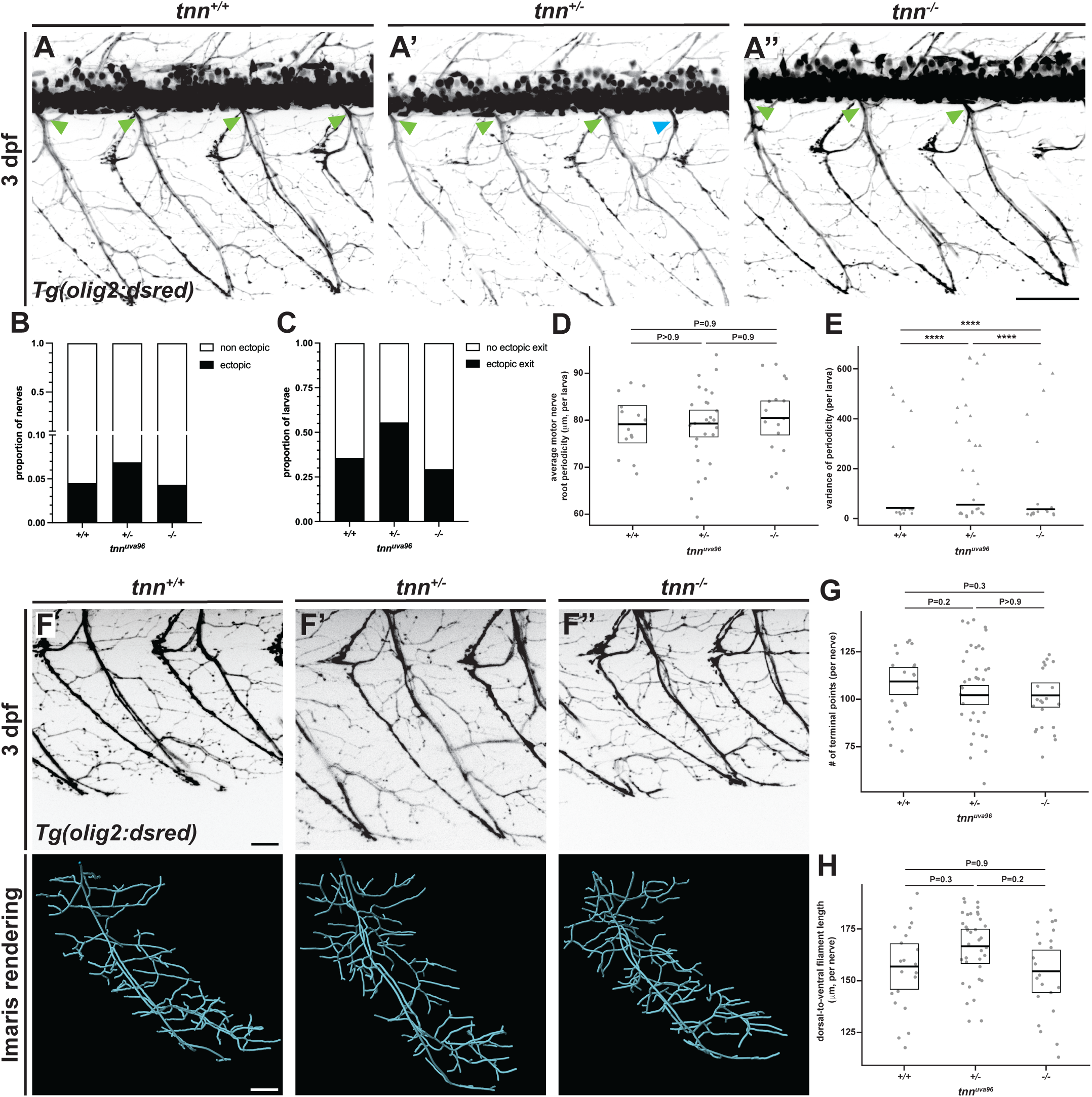
Disruptions to peripheral spinal motor nerve exit and arborization are transient in *tnn^uva96^*homozygous larvae. (**A**) Representative images from *in vivo,* confocal images of laterally mounted *Tg(olig2:dsred)* wild-type (**A**), *tnn* heterozygous (**A’**), and *tnn* homozygous (**A’’**) larvae at 3 dpf showing instances of normally placed ventral motor nerve exits (green arrowheads) and ectopic motor nerve exits (blue arrowheads). (**B**) Stacked histogram of proportion of nerves that resulted from ectopic exit and non-ectopic exit (wild-type n=155, heterozygous n=305, homozygous n=185; Fisher’s exact test P=0.4). (**C**) Stacked histogram of proportion of larvae with at least one instance of ectopic nerve exit and larvae with no instances of ectopic exit (wild-type: n=14, heterozygous: n=27, homozygous: n=17; Fisher’s exact test P=0.2). (**D**) Scatter plot of average periodicity per larva measured as distance between each nerve exit and its nearest neighbor (dot = 1 larva; wild-type n=14, heterozygous n=27, homozygous n=17). (**E**) Scatter plot of average variance of periodicity per larva (dot/triangle = 1 larva; triangle = larva with at least one instance of ectopic exit; wild-type n=14, heterozygous n=27, homozygous n=17). **** (P<0.0001) (**F**) Representative images (top images) from *in vivo* confocal imaging of laterally mounted *Tg(olig2:dsred)* wild-type (**F**), *tnn* heterozygous (**F’**), and *tnn* homozygous (**F’’**) larvae at 3 dpf enlarged to show a single nerve selected for tracing with the corresponding Imaris 3D rendering produced from tracing shown below. (**G**) Scatter plot of the number of terminal points per nerve (dot = 1 nerve; wild-type n=21, heterozygous n=36, homozygous n=22). (**H**) Scatter plot of the distance from the most dorsal point to most ventral point of each traced nerve (dot = 1 nerve; wild-type n=21, heterozygous n=36, homozygous n=22). Summary statistics for each scatter plot represent the emm with 95% CI predicted from the described model. P-values generated from post-hoc Tukey HSD. All images oriented with anterior to the left, dorsal to the top. All images taken between somites 10 and 22. Scale bars: (**A’’**) 50 μm; (**F**) 20 μm.

### *tnc* and *tnn* control distinct aspects of spinal motor nerve development

Because *tnc* is known to play a role in spinal motor nerve development in zebrafish, we hypothesized that *tnc* and *tnn* may play synergistic roles in this process. Therefore, we designed and created a gRNA to target *tnc,* creating mosaic knockdowns (termed crispants), in our *tnn* mutant larvae to test whether loss of both genes resulted in persistence of increased incidence of ectopic motor nerve exits in homozygous mutant larvae at 3 dpf. To do this, we designed a gRNA targeting the eighth exon of the *tncb* gene (Figure 6A-B). We injected this gRNA with Cas9 protein into *tnn* heterozygous and homozygous mutant embryos with *Tg(olig2:dsred)* in the background at the one-cell stage. We raised these embryos to 3 dpf and performed the same *in vivo* imaging protocol described above, followed by confirmation of cutting efficiency by Sanger sequencing (Figure 6C). Of all nerves imaged, 7.1% were the result of ectopic exit in *tnn* heterozygous/*tnc* crispant larvae compared to 2.0% in *tnn* homozygous/*tnc* crispant larvae (Figure 6D; Fisher’s exact test P=0.1). Further, 66.7% percent of *tnn* heterozygous/*tnc* crispant larvae displayed at least one ectopic nerve exit compared to 22.2% of *tnn* homozygous/*tnc* crispant larvae (Figure 6E; Fisher’s exact test, P=0.08). The average periodicity in *tnn* heterozygous/*tnc* crispant larvae was decreased compared to *tnn* homozygous/*tnc* crispant larvae (P=0.1) (*tnn* heterozygous/*tnc* crispant emm [95% CI]=75.2 [71.0, 79.4], *tnn* homozygous/*tnc* crispant emm [95% CI]=80.5 [75.7, 85.4]) (Figure 6F). The variance of periodicity was significantly increased in *tnn* heterozygous/*tnc* crispants compared to *tnn* homozygous/*tnc* crispants (P<0.0001) (*tnn* heterozygous/*tnc* crispant emm [95% CI]=95.0 [94.3, 95.8] *tnn* homozygous/*tnc* crispant emm [95% CI]=40.1 [39.7, 40.6]) (Figure 6G). These findings replicate the same trends we observed when comparing uninjected heterozygous and homozygous *tnn* mutants at 3 dpf, leading us to conclude that *tnc* does not contribute to motor axon exit at this timepoint via genetic compensation in *tnn* homozygous mutants, though it may be contributing to this phenotype independently.

**Figure 6.**
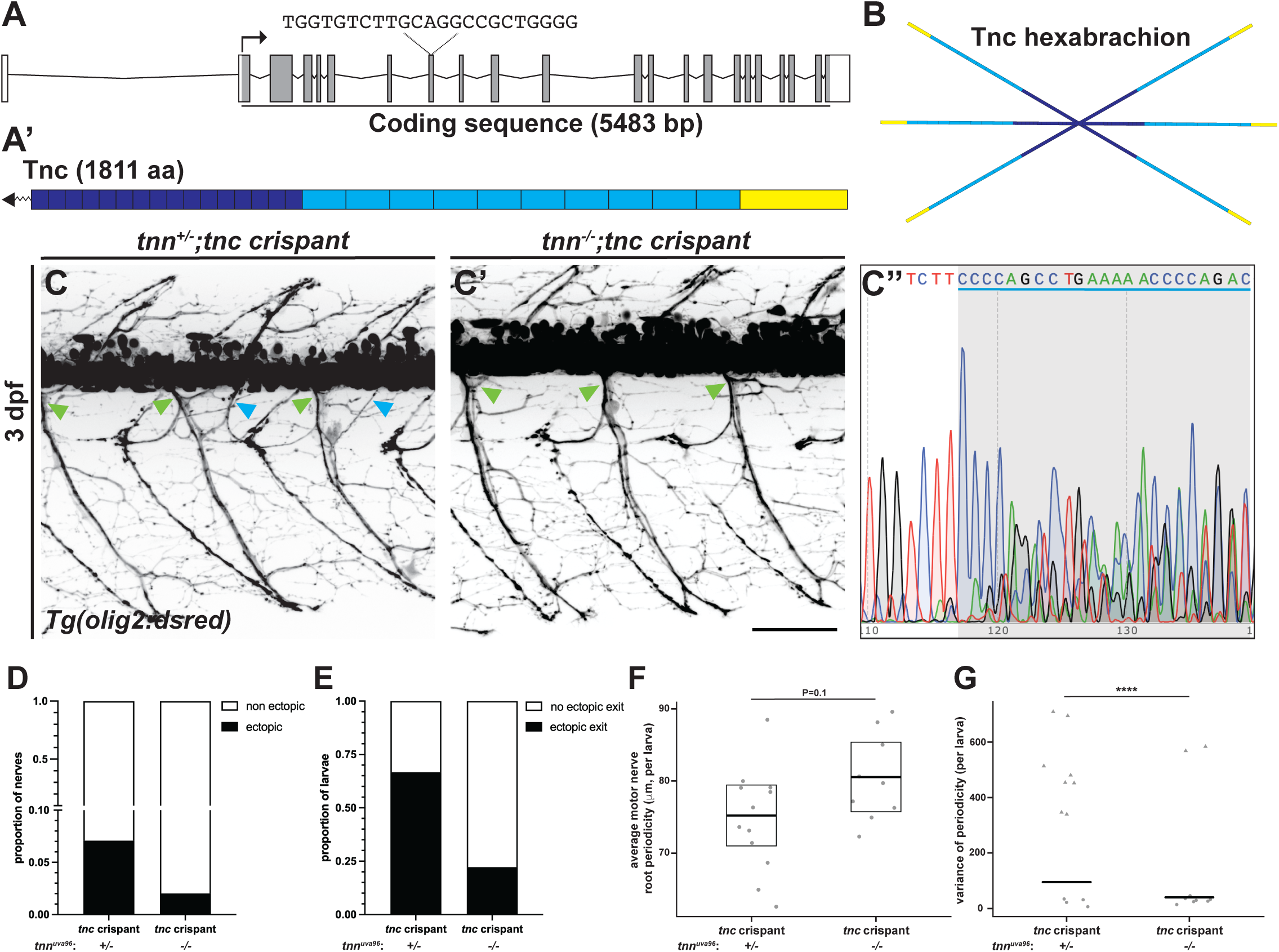
*tnc* and *tnn* control distinct aspects of spinal motor nerve development. (**A**) Schematic of zebrafish *tncb* gene. The coding sequence (grey; 5483 bp) encodes the protein domains represented in (**A’**), and the non-coding sequence is represented in white. Dashed lines above exon 8 in schematic indicate the gRNA sequence and target region of the coding sequence. (**A’**) Schematic of Tnc protein made in wild-type (1811 aa; dark blue: epidermal growth factor-like domains, light blue: fibronectin type III domains, yellow: fibrinogen-related domain). (**B**) Schematic showing that wild-type Tnc assembles into a hexabrachion made up of two Tnc trimers. (**C**) Representative images from *in vivo,* confocal images of laterally mounted *Tg(olig2:dsred) tnn* heterozygous/*tnc* crispant (**C**) and *tnn* homozygous/*tnc* crispant larvae (**C’**) larvae at 3 dpf showing instances of normally placed ventral motor nerve exits (green arrowheads) and ectopic motor nerve exits (blue arrowheads). (**C’’**) Example chromatogram from a *tnc* crispant larvae showing multiple overlapping peaks after the target sequence of the gRNA (blue underline, grey shaded region) representing multiple different alleles generated from gRNA/Cas9 activity. (**D**) Stacked histogram of proportion of nerves that resulted from ectopic exit and non-ectopic exit (*tnn* heterozygous/*tnc* crispant n=127, *tnn* homozygous/*tnc* crispant n=99; Fisher’s exact test P=0.12). (**E**) Stacked histogram of proportion of larvae with at least one instance of ectopic nerve exit and larvae with no instances of ectopic exit (*tnn* heterozygous/*tnc* crispant n=12, *tnn* homozygous/*tnc* crispant n=9; Fisher’s exact test P=0.081). (**F**) Scatter plot of average periodicity per larva (measured as distance between each nerve exit and its nearest neighbor) (dot = 1 larva; *tnn* heterozygous/*tnc* crispant n=12, *tnn* homozygous, *tnc* crispant n=9). (**G**) Scatter plot of average variance of periodicity per larva (dot/triangle = 1 larva; triangle = larva with at least one instance of ectopic exit; *tnn* heterozygous/*tnc* crispant n=12, *tnn* homozygous/*tnc* crispant n=9). **** (P<0.0001) Summary statistics for each scatter plot represent the emm with 95% CI predicted from the described model. P-values generated from post-hoc Tukey HSD. Images oriented with anterior to the left, dorsal to the top. Images taken between somites 10 and 22. Scale bar: (**C’**): 50 μm. Chromatogram generated from SnapGene.

### CRISPR/Cas9 mosaic mutagenesis of *tnn* does not affect spinal motor nerve exit or motor axon arborization at 3 dpf

Previous studies demonstrate that in stable zebrafish mutant lines, models of known disease-causing mutations sometimes do not recapitulate previously reported phenotypes due to genetic compensation (Kok et al., 2015; Rossi et al., 2015). However, transient knockdown approaches or creating mosaic mutants via injection of a gRNA and Cas9 mRNA (or protein) at the one-cell stage can mitigate the effects of genetic compensation, and this method can result in gene-specific phenotypes (Moreno et al., 2018; She, et al., 2019; Buglo, et al., 2020). To investigate whether broad genetic compensation of *tnn* contributed to the phenotypic differences we observed at 2 versus 3 dpf in *tnn^uva96^*mutant larvae, we utilized the CRISPR/Cas9 transient knockdown approach again. We injected the *tnn-*targeted gRNA described above and Cas9 protein (or Cas9 alone as a control) into *Tg(olig2:dsred)* embryos at the one-cell stage creating mosaic mutants. We raised *tnn* crispant and Cas9-only injected larvae to 3 dpf and used *in vivo* imaging to characterize motor nerve exit and branching, followed by Sanger sequencing to confirm cutting efficiency (Figure 7A). In these studies, we did not observe any differences between Cas9-injected and *tnn* crispant larvae in proportion of ectopic nerves (Cas9-injected=2.6%, *tnn* crispant=1.7%; Fisher’s exact test, P=0.5) nor proportion of larvae with at least one ectopic nerve exit (Cas9-injected=27.8%, *tnn* crispant=21.1%; Fisher’s exact test, P=0.7) (Figure 7B,C). Additionally, there were no differences in average periodicity per larvae (P=0.8) (Cas9-injeceted emm [95% CI]=81.5 μm [79.1, 84.0], *tnn* crispant emm [95% CI]=81.1 μm [78.7, 83.5]), but variance of periodicity was significantly increased in *tnn* crispants (Cas9-injected emm [95% CI]=34.4 [34.0, 34.8], *tnn* crispant emm [95% CI]=43.1 [42.4, 43.9]; P <0.0001) (Figure7 D,E). Additionally, we performed the same tracing and analysis of the ventral portion of peripheral motor nerves as described above (Figure 7F). We observed no significant differences in motor axon arborization measured by terminal points in *tnn* crispants compared to control larvae (P=0.5) (Cas9-injected emm [95% CI]=75.1 [67.9,83.0], *tnn* crispant emm [95% CI]=78.5 [70.6,87.4] terminal points) (Figure 7G-H). These findings, taken together, lead us to conclude that genetic compensation is unlikely the sole mechanism driving the differences in the phenotypes between 2 and 3 dpf in the stable knockout of *tnn*.

**Figure 7.**
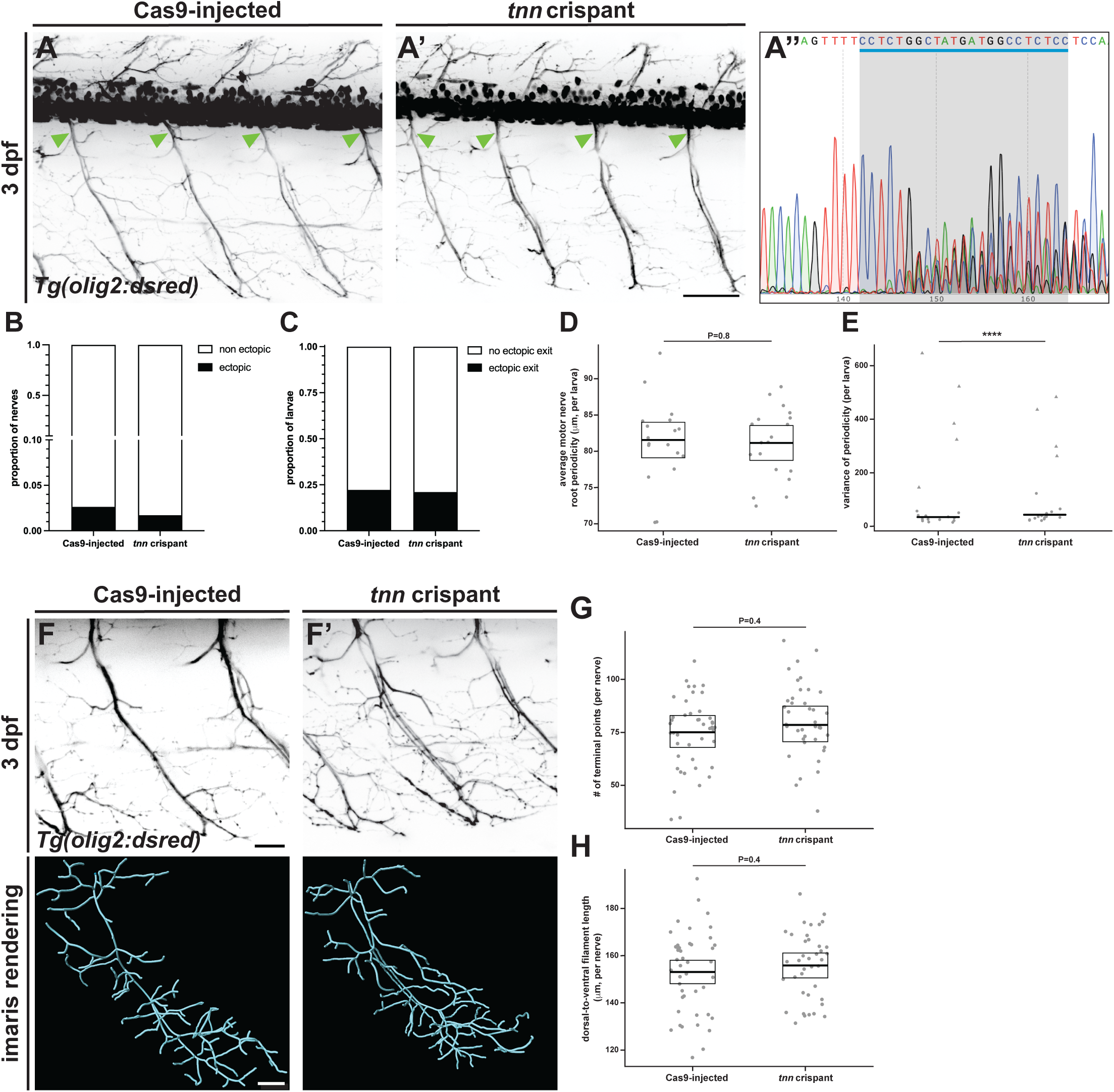
CRISPR/Cas9 mosaic mutagenesis of *tnn* does not affect peripheral spinal motor nerve exit or motor axon arborization at 3 dpf. (**A**) Representative images from *in vivo,* confocal images of laterally mounted *Tg(olig2:dsred)* Cas9-injected control (**A**) and *tnn* crispant larvae (**A’**) at 3 dpf showing instances of normally placed ventral motor nerve exits (green arrowheads). (**A’’**) Example chromatogram from a *tnn* crispant larvae showing multiple superimposed sequences after the target sequence of the gRNA (blue underline, grey shaded region). (**B**) Stacked histogram of proportion of nerves that resulted from ectopic exit and non-ectopic exit (Cas9 control n=228, *tnn* crispant n=234; Fisher’s exact test P=0.5). (**C**) Stacked histogram of proportion of larvae with at least one instance of ectopic nerve exit and larvae with no instances of ectopic exit (Cas9 control n=18, *tnn* crispant n=19; Fisher’s exact test P=0.7). (**D**) Scatter plot of average periodicity per larva (measured as distance between each nerve exit and its nearest neighbor) (dot = 1 larvae; Cas9 control n=18, *tnn* crispant n=19).. (**E**) Scatter plot of average variance of periodicity per larva (dot/triangle = 1 larva; triangle = larva with at least one instance of ectopic exit; Cas9 control n=18, *tnn* crispant n=19). **** (P<0.0001) (**F**) Representative images (top images) from *in vivo* confocal imaging of laterally mounted *Tg(olig2:dsred)* Cas9 control (**F**) and *tnn* crispant (**F’**) larvae at 3 dpf enlarged to show a single nerve selected for tracing with the corresponding Imaris 3D rendering produced from tracing shown below. (**G**) Scatter plot of the number of terminal points per nerve (dot = 1 nerve; Cas9 control n=44, *tnn* crispant n=39). (**H**) Scatter plot of the distance from the most dorsal point to most ventral point of each traced nerve (dot = 1 nerve; Cas9 control n=44, *tnn* crispant n=39). Summary statistics for each scatter plot represent the emm with 95% CI predicted from the described model. P-values generated from post-hoc Tukey HSD. All images oriented with anterior to the left, dorsal to the top. All images taken between somites 10 and 22. Scale bars: (**A’**) 50 μm; (**F**) 20 μm. Chromatogram generated from SnapGene.

## Discussion

In this study, we investigated the role of an understudied ECM protein, Tnn, in the development of spinal motor nerves. While we know from previous studies that the ECM plays important roles throughout this process, many of the proteins and their specific roles are still unknown. Here, we describe a subtle and transient role for Tnn in motor axon exit and arborization in developing zebrafish.

*Tnn* is expressed as early as 24 hpf in zebrafish embryos in cells adjacent to the neural tube and outgrowing motor axons (Weber et al., 1998). However, it was not known whether *tnn* contributed to development of spinal motor nerves. We show that *tnn*/Tnn is expressed and localized along vertical myosepta and the border of the ventral neural tube from 1 to 3 dpf, the window in which motor neurons extend axons out of the spinal cord and grow toward their muscle targets. Additionally, we created a novel null mutant allele, *tnn^uva96^,* to interrogate the function of *tnn* during this critical window of peripheral nervous system development. With *in vivo* imaging and morphological analysis, we show that loss of *tnn* results in a subtle and transient increased incidence of ectopic motor axon exit from the neural tube and increased arborization of motor axons at 2 dpf. However, by 3 dpf, only heterozygous mutant larvae show increased incidence of ectopic nerve exits and a significantly increased variance of periodicity, leading us to conclude that *tnn* contributes to early spinal motor development, but that loss of two copies of *tnn* may induce a genetic compensation response.

The appearance of ectopic motor axons is described in multiple studies, often resulting from loss of diffusible guidance cues and/or their cognate receptors (Birely et al., 2005; Palaisa & Granato, 2007; Feldner, et al., 2007). In these studies, ectopic exits were commonly caused by aberrant placement of primary motor neurons or misguided axons from primary motor neurons to MEP TZs (Birely et al., 2005; Palaisa & Granato, 2007; Feldner, et al., 2007). In our study, we did not observe expression of *tnn/*Tnn within the neural tube nor did we see any ectopic exits until after primary motor neurons had already exited the spinal cord, which leads us to hypothesize that *tnn* does not impact primary motor neuron cell body placement nor guidance of primary motor axons out of the spinal cord. The localization of Tnn adjacent to the ventral neural tube from 2 dpf on and increased incidence of ectopic exits at this timepoint lead us to hypothesize that *tnn* loss affects permissibility of axonal exit from the spinal cord. This could be a consequence of a direct mechanism that restricts axonal exit, where growing neurites contact Tnn at the boundary of the neural tube through a cell surface receptor, or an indirect mechanism, by which the loss of *tnn* could result in alterations to the ECM architecture surrounding the neural tube, leading to domains of increased axonal permissibility. All members of the tenascin family have predicted integrin binding sites within their FNIII domains supporting a possible direct mechanism (Chiquet-Ehrismann & Tucker, 2011; Tucker & Degen, 2019). For example, Neidhardt and colleagues showed that a substrate coated with a small splice variant of TNN repelled hippocampal neurites *in vitro*, and Degen and colleagues demonstrated that fibroblast cells adhere to a human TNN-coated substratum in a β1-integrin dependent manner (Neidhardt et al., 2003; Degen et al., 2007). Recent findings of Nemoz-Billet and colleagues, however, offer support for an indirect mechanism due to ECM rearrangement (Nemoz-Billet et al., 2024). They demonstrated that loss of *col15a1b* results in the disorganization of Tnc in the path of growing motor axons, and this ultimately leads to increased branching of motor axons at 27 hpf, a finding they also observe with the loss of *tnc* (Nemoz-Billet et al., 2024). Deciphering whether the loss of *tnn* leaves growing axons without a source of contact or changes the overall ECM architecture would provide a clearer understanding by which increased ectopic exits occur. While both hypotheses are possible causes of the ectopic phenotype we observe, we hypothesize that an indirect mechanism is more likely the cause of the increased branching we saw, primarily because Tnn is not localized in the pathway of growing motor axons. Therefore, it seems more likely that the loss of Tnn along vertical myosepta could cause reorganization of adjacent ECM proteins, creating an environment more permissible to branching. Uncovering which specific cell types are responsible for *tnn* expression along vertical myosepta and the ventral neural tube will aid in our understanding of which of these mechanisms drives the changes we observed.

Although we showed a similar pattern of expression and localization at 2 and 3 dpf, we observed a change in phenotypes between heterozygous and homozygous mutant larvae at 3 dpf. The increased incidence of ectopic exit did not persist at 3 dpf in homozygous mutant larvae but did in heterozygotes. Similarly at 3 dpf, *tnn* heterozygous larvae with *tnc* knockdown showed an even larger increased incidence of ectopic exit compared to *tnn* homozygous larvae with *tnc* knockdown. Lastly, mosaic knockdown of *tnn* in wild-type embryos did not result in persistence of increased ectopic exits nor arborization at 3 dpf. Additionally, at both 1 and 3 dpf despite non-significant changes to incidence of ectopic exit and periodicity, *tnn* heterozygotes displayed significant increases in variance of periodicity compared to wild-types and homozygotes, further suggesting *tnn* has subtle impacts on the spacing of spinal motor axon exit.

All of this together, led us to two major conclusions. First, the loss of two copies of *tnn* may induce a more robust genetic compensation than the loss of just one. Recent studies have begun to define the mechanisms underlying the genetic compensation response (Ma et al., 2019; El-Brolosy et al., 2019). These efforts show that mutant mRNA containing a premature termination codon and nonsense-mediated decay proteins trigger the genetic compensation response, and that for some genes this effect is less robust in heterozygous mutants than homozygous (Ma et al., 2019; El-Brolosy et al., 2019). This could be the case for loss of *tnn* as we observed reduced, but not absent, expression of *tnn* transcripts in heterozygous larvae compared to wild-type larvae via RNA *in situ* hybridization assays (data not shown). The genetic compensation response in both heterozygous and homozygous larvae could also be contributing to the subtlety of the phenotypes we observed, which leads into our second conclusion. Our results highlight the complexity of the network of proteins that contribute to the assembly of spinal motor nerves. Transient phenotypes during this critical window of spinal motor nerve development have also been reported for Tnc and cannot be attributed to genetic compensation alone (Schweitzer et al., 2005). For example, overexpressing of either the Tnc protein or RNA encoding TNC’s EGF domains resulted in truncated motor axon extension at 24 hpf but resolved by 33 hpf (Schweitzer et al., 2005). Further, reduction of *tnc* expression resulted in aberrant branching not seen at 24 hpf but was apparent by 33 hpf (Schweitzer et al., 2005). Notably, these phenotypes were not investigated beyond 2 dpf. While much of our data here show trends of increased incidence of ectopic exit and motor axon arborization that lack statistical significance, we do not believe that this diminishes that Tnn functions in these processes. We believe it only adds to the already established conclusion that the ECM network that orchestrates the assembly of peripheral motor nerves is vast and complex, and that Tnn is just one molecular mediator of many. The loss of Tnn may not be detrimental to the development of peripheral motor nerves, but it’s loss certainly results in changes to the process.

In conclusion, we describe a novel role for an understudied ECM protein, Tnn, in early spinal motor nerve morphogenesis. Our findings open the door to further investigation of the mechanisms by which Tnn affects this process and how it fits into the broader network of ECM proteins that influence peripheral motor nerve development. Our work further emphasizes the need to continue deciphering the connections in this complex network and how all of these proteins work in concert to achieve construction of peripheral motor nerves

## Materials & Methods

### Zebrafish Husbandry

All animal studies were approved by the University of Virginia Institutional Animal Care and Use Committee (IACUC). Animals were housed at 28°C at a density of 8 to 10 fish/liter and exposed to a 14/10 hour light-dark cycle. Previously published zebrafish lines used in this study are included in section “*Zebrafish transgenic lines.*” Newly created zebrafish lines from this study are described in “*Generation of stable tnn mutant line.*” Embryos were produced by pairwise, natural matings and raised in egg water at 28.5°C. All embryos and larvae were staged according to hours or days post fertilization (Kimmel et al., 1995) and screened for desired transgenic fluorescence prior to use. Sex was not considered as a variable for experiments because sex is indeterminate in zebrafish prior to 25 dpf (Takahashi, 1977).

### Zebrafish transgenic lines

Previously published transgenic lines used in this study include *Tg(olig2:dsred2)^vu19^*(Shin et al., 2003), and *Tg(nkx2.2a:meGFP)^vu17^* (Kucenas et al., 2008b). All transgenic lines described are stable and incorporated into the germline.

### Guide RNA (gRNA) template generation and transcription

We identified *tnn* and *tncb* gRNAs using *CRISPRscan* (Moreno-Mateos et al., 2015). We synthesized these templates through PCR amplification with a gene specific forward primer and the following constant oligonucleotide reverse primer, 5′-*AAAAGCACCGACTCGGTGCCACTTTTTCAAGTTGATAACGGACTAGCCTTATTTTAACTTGCTATTTCTAGCTCTAAAAC*-3′. For the *tnn* gRNA, we targeted 5’-*CCTCTGGGCCTATAAGGGCTCTC*-3’ in the sixth coding exon and synthesized it using the following forward primer: 5’-*taatacgactcactataGGGAGCCCTTATAGGCCCAGgttttagagctagaaATAGCAAG*-3’. For the *tncb* gRNA, we targeted 5’-*TGGTGTCTTGCAGGCCGCTGGGG*-3’ in the eighth exon and synthesized it using the following forward primer 5’-*taatacgactcactataGGGTGTCTTGCAGGCCGCTGgttttagagctagaaATAGCAAG*-3’. To synthesize gRNA templates, we conducted PCR, gel electrophoresis to confirm product size, and purification using QIAquick PCR purification kit (Qiagen). After purification, gRNAs were transcribed using Megascript (AMBION) or HiScribe T7 RNA synthesis kit (New England Biolabs) and precipitated via lithium chloride precipitation.

### Generation of stable *tnn* mutant line and crispants

To create a *tnn* mutant allele, *tnn* crispants, and *tnc* crispants, we injected a mixture containing gRNA (200 ng/μl) and Cas9 protein into embryos at the one-cell stage (or just Cas9 protein alone for Cas9 controls). For the stable *tnn* mutant line, we confirmed insertion and deletion mutations (INDELs) by extracting genomic DNA via HotSHOT (hot sodium hydroxide and tris), followed by PCR amplification using the following primers: forward primer, 5′-*TGACGGAAAACTCTGCGACT*-3′, and reverse primer, 5′-*TCCCTTGAGGAAACGCTTGA*-3′. Purified DNA was then sent for Sanger sequencing through GENEWIZ (Azenta Life Sciences) to confirm successful mutagenesis (Truett et al., 2000). We identified founders and confirmed germline integration by outcrossing F_0_ adults to WT adults and analysis of genomic DNA of F_1_ offspring via Sanger sequencing. We first identified mutant alleles in F_1_ adults via analysis of Sanger sequencing. To confirm suspected mutant alleles, we cloned the PCR product into pCR4-TOPO TA vector (Invitrogen), amplified vectors containing *tnn* mutant alleles in chemically competent E. coli, isolated DNA via miniprep (Qiagen), and analyzed Sanger sequencing for INDELs. We identified F_1_ adults containing a 4 base-pair insertion in the *tnn* gene, and these were raised to establish a stable mutant line for further experimentation (*tnn^uva96^)*. All subsequent generations were genotyped as described below in *“Genotyping”*.

### Genotyping and crispant confirmation

Genomic DNA was extracted using HotSHOT (hot sodium hyrodxide and tris) (Truett et al., 2000). For genotyping of the *tnn^uva96^* allele, PCR amplification was performed using primers listed above, followed by restriction enzyme digestion with Sau96I restriction endonuclease (New England Biolabs). For the digestion, samples were incubated for 1 hour at 37°C followed by heat inactivation for 20 minutes at 65°C. The resulting digest products were screened with gel electrophoresis on a 2% agarose gel.

To confirm mosaic mutagenesis in *tnn* and *tncb* crispants, larvae were recovered from imaging dishes after experimentation, and the same procedures were used as described above, except: following PCR amplification, PCR products were purified using QIAquick PCR purification kit (Qiagen) and sent for sequencing through GENEWIZ (Azenta Life Sciences). The resulting chromatograms were analyzed for effective Cas9 cutting, and data acquired from larvae displaying only wild-type sequence was discarded. Primers used for *tncb* amplification were as follows: forward primer 5’-*TCTGGATTCCCCAAAGCACC*-3’ and reverse primer 5’-*TCTCCAATGTCTGGTTCGCA*-3’.

### In vivo Imaging

Embryos were treated with 0.003% PTU at 24 hpf to reduce pigment formation and screened for transgenic fluorescence prior to imaging. For stages before 2 dpf, embryos were manually dechorionated. Embryos and larvae were immobilized prior to imaging using diluted Tricaine (MS-222; Syndel/Western Chemical, ordered from The Pond Outlet; concentration varied by age). Larvae were then mounted laterally in 0.8% low gelling point agarose in 4-well glass bottom 35 mm Petri dishes (Fisher, Greiner Bio-One) and covered with diluted Tricaine (concentration varied by age), unless otherwise noted. Imaging was performed on a motorized Zeiss AxioObserver Z1 microscope equipped with a Quorum WaveFX-XI (Quorum Technologies), Andor CSU-W1 Spinning Disc Confocal Microscope, or Andor Dragonfly 200 Spinning Disc Confocal Microscope with iXon Ultra camera with 40x/1.10 objective (Andor/Oxford Instruments). Images were illuminated with 488 nm (GFP, mEGFP) and 561 nm (RFP, dsRed) lasers. For still imaging of spinal motor nerves, three images were captured starting at the level of the 10^th^ somite and moving posteriorly along the spinal cord. Time-lapse imaging also occurred within this three-field-window beginning at the 10^th^ somite. Time-lapse length and time-step varied according to the feasibility and needs of the experiment and is stated if relevant. Images were acquired as 60 μm z-series with 2 μm step size unless otherwise stated.

### Cryosectioning

Fixed larvae were mounted in sectioning agar (1.5% agar; Fisher Scientific; 5% sucrose; Sigma; 100 mL ultrapure water) and submerged in 30% sucrose in 1x phosphate buffered saline (PBS) overnight at 4°C. Agar blocks were frozen by placing them on a small raft floating in 2-methylbutane (Fisher Scientific), the container of which was submerged in a bath of liquid nitrogen. Blocks were sectioned to a thickness of 20 or 40 µm (depending on experiment) with a cryostat microtome and mounted on microscope slides (VWR). Slides were air-dried at room temperature for 3 hours and then stored at –20°C or –80°C until needed for immunohistochemistry (IHC) or *in situ* hybridization (ISH). Prior to sectioning for IHC or ISH experiments performed on *tnn^uva96^*WT, heterozygote, and mutant clutchmates, larvae were anesthetized in diluted Tricaine (MS-222; Syndel/Western Chemical, ordered from The Pond Outlet; concentration varied by age), and the heads were removed with a scalpel. Heads were kept for genotyping, and the trunks were fixed for sectioning.

### Wholemount immunohistochemistry

Embryos and larvae (1 to 3 dpf) were dechorionated (if necessary), fixed in 4% paraformaldehyde (PFA; Sigma) for 2 hr shaking at room temperature (RT) for all experiments, and used immediately after for antibody staining or stored in 100% methanol for future use. Embryos/larvae were washed in PBSTx (1% Triton X-100, 1x PBS) 3 x 10 minutes shakings and then incubated with Proteinase K (diluted in PBSTx to final concentration of 20 μg/ml) for 20 minutes at RT. Embryos/larvae were then washed 2 x 5 minutes shaking in PBSTx, fixed again in 4% PFA (diluted in PBSTx) shaking for 20 minutes at RT, and washed 3 x 5 minutes shaking at RT. Embryos/larvae were then blocked in 5% goat serum + 1% DMSO in PBSTx for 1-2 hours shaking at at RT, and then overnight at 4°C with antibodies in blocking solution. After antibody incubation, embryos/larvae were washed in PBSTx 4 x 10 minutes shaking at RT and incubated in secondary antibody for 2 hours shaking at RT or overnight at 4°C. Embryos/larvae were washed extensively with PBSTx and stored in 50% glycerol/50% PBS at 4°C until imaging. The following antibodies were used: rabbit anti-Tenascin-N (1:500; MyBioSource), mouse anti-acetylated tubulin (1:10,000; Sigma), Alexa Fluor 488 goat anti-rabbit IgG(H+L) (1:400; Thermo Fisher), and Alexa Fluor 568 goat anti-mouse (1:400; Thermo Fisher). Fish were immobilized and imaged in glass-bottom dishes with microscopes described in “*In vivo imaging”*.

### Immunohistochemistry on section

Embryos were fixed in 4% PFA for 2 hours at RT and sectioned as described in “*Cryosectioning”.* Sections were rehydrated in 1X PBS for 10 minutes and blocked with 2% goat serum + 2 mg/ml bovine serum albumin (BSA; Fisher Scientific) in PBS for 30 minutes to 1 hour at RT. Sections were incubated in primary antibody overnight at 4°C. Then, sections were washed continuously with PBS for 45 minutes followed by secondary antibody incubation for 2 hours at RT. The same antibodies and concentrations were used as described in “*Wholemount immunohistochemistry*”.

Slides were stored in the dark until they were imaged using microscopes described in “*In vivo imaging”*.

### In situ probe synthesis

The *tnn* RNA probe for chromogenic ISH was designed and synthesized in our lab as described in Fontenas and Kucenas, 2021. The primers used to generate the *tnn* RNA probe were as follows: forward primer, 5’-*AGGAATCAAGGTGCTGAGCC*-3’, and reverse primer plus T7 sequence, 5’-*TAATACGACTCACTATAGAGCTGGTAAGATGTGGCACC*-3’. For fluorescent in situ hybridization experiments (FISH), the *tnn* RNA probe (RNAscope probe Dr-tnn-C3) was purchased from Advanced Cell Diagnostics (ACD).

### Chromogenic *in situ* hybridization

Embryos and larvae (1 to 3 dpf) were dechorionated, if necessary, and fixed in 4% PFA for 2 hr shaking at RT and then transferred to 100% MeOH overnight at –20°C. Chromogenic ISH was performed as described in Hauptmann and Gerster, 2000. Images were obtained either using a Zeiss AxioObserver inverted microscope equipped with Zen, using a ×40 oil immersion objective, or a Zeiss Axiozoom V16 microscope.

### Fluorescent *in situ* hybridization (RNAscope)

Larvae (1 to 3 dpf) were dechorionated, if necessary, and fixed with 4% PFA for 2 hr shaking at RT, dehydrated in 100% MeOH overnight at –20°C, and cryosectioned (see ‘*Cryosectioning*’). RNAscope was performed as described in Wiltbank et al., 2022. Sections were imaged using equipment described in “*In vivo imaging”*.

### RT-PCR analysis

RNA was extracted using TRIzol reagent (Invitrogen) from 2 dpf *tnn^uva96^*and AB* embryos. cDNA libraries were generated from the RNA using the high-capacity cDNA reverse transcription kit (Thermo Fisher). Using the cDNA as a template, PCR amplification was performed with the following primers: forward primer, 5′-*GCCAGGGAGGACATGAGAAC*-3′, reverse primer, 5′-*ACAGTCGCAGAGTTTTCCGT*-3′. The resulting PCR products were visualized with gel electrophoresis on a 2.5% agarose gel.

### Motor nerve periodicity measurements

*In vivo* images were processed using FIJI. Images were max projected (and rotated if necessary), and then the multi-point tool was used to mark each motor nerve exit from the spinal cord. After marking all exits, the “measure” function was selected and the “X” value was exported to excel for each image. Periodicity was calculated by subtracting each “X” value (location of a nerve exit) from the next closest “X” value in the posterior direction within each image. Suspected ectopic exits were confirmed by examining single slices in images stacks in FIJI.

### Imaris filament tracing

Nerve branching was quantified using Imaris (Oxford Instruments) first by cropping images containing multiple nerves and the spinal cord to include only the portion of a single nerve and its constituent branches ventral to the spinal cord. Then, we used the filament tracer module in the Imaris software to semi-automatically trace the nerve and its axonal branches, followed by manual corrections and additions missed by the semi-automatic function. In the resulting statistics, we selected segment terminal points (a count of each axonal ending) as a measurement for axonal branching. After tracing of each nerve was completed, we used the measurement points module in Imaris to measure the distance of the most dorsal point to the most ventral point of the trace mask. This measurement was then included in the statistical analysis of terminal points (see quantification and statistical analysis).

### Quantification and statistical analysis

We conducted quantitative analyses to assess ectopic nerve counts, periodicity of spinal motor nerve exit, and spinal motor nerve branching using Imaris, FIJI, GraphPad Prism, R (vers. 4.5.2), and R Studio Software (vers. 2023.06.1). Periodicity and nerve branching were measured as described above. Proportion of ectopic nerves and larvae with at least one instance of ectopic nerves were compared in GraphPad Prism using Fisher’s exact test. Width of the 10^th^ somite was compared in GraphPad Prism using one-way ANOVA with Tukey’s post-hoc test. Periodicity data analyzed on the nerve level via linear mixed model (LMM) acconting for random effects of larva and image. Variance of periodicity data analyzed on the inage level via gamma distribution mixed model accounting for random effect of larva. Terminal points data analyzed on the nerve level via Poisson mixed model accounting for random effecy of larva and offset for dorsal-to-ventral length. Dorsal-to-ventral trace length data analyzed on the nerve level via LMM accounting for random effects of larva. Mixed effects models to compare motor nerve periodicity, variance of periodicity, terminal points, and dorsal-to-ventral trace length were generated in R. P-values for mixed models generated from post-hoc Tukey HSD. See GitHub link (will be added after review) for code used to generate statistical analyses.

## Acknowledgements

We thank members of the Kucenas lab, past and present, for valuable discussions and training, and especially Lori Tocke for zebrafish care. This work was funded by the NIH/NIGMS (T32GM007267, CGM), Whitfield Randolph Scholarship (CGM), and NIH/NINDS (R01NS107525, SK).

## Author Contributions

Conceptualization, C.G.M. and S.K.; Methodology, C.G.M. and S.K.; Investigation, C.G.M., C.I.B., and S.K.; Writing – Original Draft, C.G.M. and S.K.; Writing – Review & Editing, C.G.M. and S.K.; Formal analysis, C.G.M., and M.J.

---

**Supplementary Video 1. Appearance of an ectopic motor axon exit in *tnn^uva96^* larvae.** A spinal motor nerve exits at an ectopic location (magenta arrowhead) between 1 and 2 dpf in a *Tg(olig2:dsred),* wild-type, heterozygous mutant, and homozygous mutant larvae. Images captured every 20 minutes over the course of 16 hours beginning between 24-28 hpf. Anterior to the left, dorsal to the top. Scale bar: 50 μm.

